# Generation and characterization of *Ccdc28b* mutant mice links the Bardet-Biedl associated gene with social behavioral phenotypes

**DOI:** 10.1101/2021.10.21.465251

**Authors:** Matías Fabregat, Sofía Niño, Sabrina Pose, Magdalena Cárdenas-Rodríguez, Corrine Smolen, Mariana Bresque, Karina Hernández, Victoria Prieto-Echagüe, Geraldine Schlapp, Martina Crispo, Patricia Lagos, Natalia Lago, Santhosh Girirajan, Carlos Escande, Florencia Irigoín, Jose L. Badano

**Author notes:** To whom correspondence should be addressed: Jose L. Badano, ^1^Institut Pasteur de Montevideo, Mataojo 2020, Montevideo CP 11400, Uruguay, Florencia Irigoín, ^8^Departamento de Histología y Embriología, Facultad de Medicina, Universidad de la República, Gral. Flores 2125, Montevideo CP11800, Uruguay; ^1^Institut Pasteur de Montevideo, Mataojo 2020, Montevideo CP 11400, Uruguay.

## Abstract

CCDC28B (coiled-coil domain-containing protein 28B) was identified as a modifier in the ciliopathy Bardet-Biedl syndrome (BBS). Our previous work in cells and zebrafish showed that CCDC28B plays a role regulating cilia length in a mechanism that is not completely understood. Here we report the generation of a *Ccdc28b* mutant mouse using CRISPR/Cas9 (*Ccdc28b mut*). After confirming the depletion of *Ccdc28b* we performed a phenotypic characterization showing that *Ccdc28b mut* animals present a mild phenotype: i) do not present clear structural cilia affectation, although we did observe mild defects in cilia density and cilia length in some tissues, ii) reproduce normally, and iii) do not develop retinal degeneration or obesity, two hallmark features of reported BBS murine models. In contrast, *Ccdc28b mut* mice did show clear social interaction defects as well as stereotypical behaviors suggestive of autism spectrum disorder (ASD). This finding is indeed relevant regarding CCDC28B as a modifier of BBS since behavioral phenotypes have been documented in BBS. Importantly however, our data suggests a possible causal link between CCDC28B and ASD-like phenotypes that exceeds the context of BBS: filtering for rare deleterious variants, we found *CCDC28B* mutations in eight probands from the Simmons Simplex Collection cohort. Furthermore, a phenotypic analysis showed that *CCDC28B* mutation carriers present lower BMI and mild communication defects compared to a randomly selected sample of SSC probands. Thus, our results suggest that mutations in *CCDC28B* lead to mild autism-like features in mice and humans. Overall, this work reports a novel mouse model that will be key to continue evaluating genetic interactions in BBS, deciphering the contribution of CCDC28B to modulate the presentation of BBS phenotypes. In addition, our data underscores a novel link between *CCDC28B* and ASD-like phenotypes, providing a novel opportunity to further our understanding of the genetic, cellular, and molecular basis of ASD.

**AUTHOR SUMMARY:** Bardet-Biedl syndrome (BBS) is caused by mutations in any of 21 genes known to date. In some families, BBS can be inherited as an oligogenic trait whereby mutations in more than one BBS gene collaborate in the presentation of the syndrome. In addition, *CCDC28B* was identified as a modifier of BBS, associated with a more severe presentation of the syndrome. Different mechanisms, all relying on functional redundancy, have been proposed to explain this genetic interaction and the characterization of different BBS proteins supported this possibility as they were shown to play roles in the same cellular organelle, the primary cilium.

Similarly, CCDC28B also participates in cilia biology regulating the length of the organelle: knockdown of CCDC28B in cells results in cilia shortening and depletion in zebrafish also results in early embryonic phenotypes characteristic of other cilia mutants. Here, we sought to generate a mouse *Ccdc28b* mutant to determine whether it would be sufficient to cause a ciliopathy phenotype, and to generate a reagent critical to further dissect its modifying role in the context of BBS. Overall, *Ccdc28b* mutant mice presented a mild phenotype, a finding fully compatible with a modifier rather than a causal BBS gene. In addition, we found that *Ccdc28b* mutants showed a clear autism-like behavior, and autism is indeed a feature of several BBS patients. Importantly, we identified multiple individuals with autism from the Simmos Simplex Collection to carry disruptive mutations in *CCDC28B* suggesting that this gene is causally associated with autism independently of BBS.

## INTRODUCTION

BBS is a rare disorder characterized by retinal degeneration, polydactyly, mental retardation, gonadal/renal malformations and obesity among other features [1]. BBS is a genetically heterogeneous condition with 21 genes known to cause the disease to date (*BBS1*-*BBS21*; [2] and references within). All the BBS proteins that have been characterized participate in the formation/maintenance of primary cilia [3–13]. Therefore, BBS is a ciliopathy, a termed used to group several human conditions that are caused by ciliary dysfunction and share, to different degrees, a set of characteristic phenotypes [14, 15]. While in most families BBS is inherited as an autosomal recessive trait, it has been shown that genetic interactions between BBS genes can modulate both the penetrance and expressivity of the syndrome, thus dubbing BBS as an oligogenic condition [16–25].

The functional characterization of BBS proteins has provided a cellular/molecular explanation to the oligogenicity observed in BBS, a phenomenon that typically relies on the presence of complementary pathways, complexes and/or some degree of functional redundancy [26]. In this context, BBS proteins present a significant functional overlap and can even interact directly forming multiprotein complexes. Eight BBS proteins form the BBSome, a complex that mediates traffic of ciliary components [5, 9, 12, 27–30]. Another group of BBS proteins (BBS6, BBS10 and BBS12) have a chaperone activity critical for BBSome assembly [31], while others are important for BBSome recruitment to membranes [5] or to regulate the movement of the complex in and out of cilia [32]. Moreover, cilia are complex organelles composed of more than 1000 proteins and with at least four main functional complexes including the BBSome, the transition zone, and two intraflagellar complexes for anterograde and retrograde transport respectively. Thus, mutations in different genes and ciliary modules can contribute to cilia dysfunction and therefore to the pathogenesis of different ciliopathies (reviewed in for example [33, 34]).

*CCDC28B* (coiled-coil domain-containing protein 28B) was originally identified as a gene associated with Bardet-Biedl syndrome (BBS; OMIM 209900) by virtue of its physical interaction with different BBS proteins [35]. Moreover, it was shown to play a modifier role in BBS, at least in some patient cohorts [18, 35–37], whereby a reduction in CCDC28B levels, in a genetic background with mutations in *bona fide* BBS genes, was shown to correlate with a more severe presentation of the syndrome [35]. We have shown that CCDC28B also plays a role in cilia, thus providing insight on the cellular basis of its genetic modifier effect. Knockdown of *CCDC28B* in hTERT-RPE cells results in shortened cilia and a reduction in the percentage of ciliated cells. Targeting *ccdc28b* in zebrafish results in a distinct external phenotype characterized by a shortened body axis, increased body curvature, craniofacial and pigmentation defects and smaller eyes, phenotypes that have been described in other cilia mutants in the fish ([38, 39] and references within). Furthermore, the analysis of different ciliated tissues in zebrafish morphant embryos showed a clear reduction in both the number and length of cilia [38–40]. While the mechanism by which CCDC28B modulates cilia is still not completely understood, we have uncovered relevant protein-protein interactions. We were able to show that CCDC28B modulates cilia length, at least in part, through an interaction with SIN1, a member of mTORC2, but independently of the mTOR complex [40]. More recently, we showed that the molecular motor kinesin 1 is also involved in cilia length regulation by controlling CCDC28B sub-cellular localization [39].

In this work we aimed to generate a *Ccdc28b* knockout mouse that would i) allow us to determine whether loss of function of this gene is sufficient to cause cilia dysfunction and associated phenotypes in mammals, and ii) serve as a tool to study its modifier effect in the context of BBS. We therefore targeted *Ccdc28b* in mice using CRISPR/Cas9 and performed an in-depth phenotypic characterization of mutant animals (*Ccdc28b mut*). We focused on phenotypes that have been described for BBS mouse mutant lines, which include the development of obesity driven by hyperphagia, and retinal degeneration. We show that depletion of Ccdc28b is not sufficient to cause overt ciliary defects in neither cells nor tissues, although a trend towards presenting a reduction in the percentage of cilia and mild cilia length defects were seen in a subset of the analyzed tissues. In agreement with this observation, *Ccdc28b mut* are viable, reproduce at mendelian rates and do not develop obesity or show signs of photoreceptor loss. *Ccdc28b mut* mice show however, autistic-like behaviors, phenotypes that are being documented in both BBS patients and animal models (see for example [41–47]). Finally, by analyzing the presence of *CCDC28B* mutations, as well as the phenotypic presentation of carrier probands, in the Simons Simplex Collection cohort we underscore a likely causal link between *CCDC28B* and mild features of autism. Our work generates a novel genetic model that recapitulates certain phenotypes observed in BBS patients, and allows for further dissection of genetic, cellular, and molecular basis of complex phenotypes relevant to both BBS and autism.

## RESULTS

### Generation of a Ccdc28b mouse model

The mouse *Ccdc28b* gene (Gene ID 66264) is located on chromosome 4 and is composed of six exons with an open reading frame spanning from exon 2 to exon 6. Although several splicing isoforms have been reported (*Ensembl* ENSMUSG00000028795), as in humans, only two transcripts are predicted to encode full length proteins in mice. These are proteins of 200 and 204 amino acids respectively, differing in their C-terminal regions (Fig 1A). To generate a knockout *Ccdc28b* murine line we choose to target exon 3, which is shared by all reported transcripts and coding isoforms, and is the exon affected by the mutation described in humans [35]. We designed two gRNAs to target the 5’ end of exon 3 (Fig 1B) and verified their efficiency in targeting *Ccdc28b* by transiently transfecting a murine NIH3T3 cell line stably expressing CAS9 and performing an heteroduplex analysis. We then injected 294 zygotes with *Cas9* mRNA and our two gRNAs. Upon analysis of 12 animals by PCR amplification of exon 3 from genomic DNA and sequencing, we were able to identify several mutations in six mice (50% mutation rate). We crossed two of those founder mice with C57BL/6J females (Jackson Lab stock # 000664) to segregate the mutations, and finally, we chose to continue our work with a mutation consisting of two one base pair deletions at the 5’ end of exon 3, thus predicted to result in a frameshift and a PTC (Fig 1B). This mutant could potentially encode an 88 amino acid protein composed of the first 56 amino acids of Ccdc28b followed by 32 novel residues, although it is expected to be degraded by nonsense-mediated decay (NMD) (Fig 1B). We crossed a male mouse carrying the selected mutation with C57BL/6J females to start the colony and performed two rounds of crossing before starting the characterization of the line.

**Figure 1:**
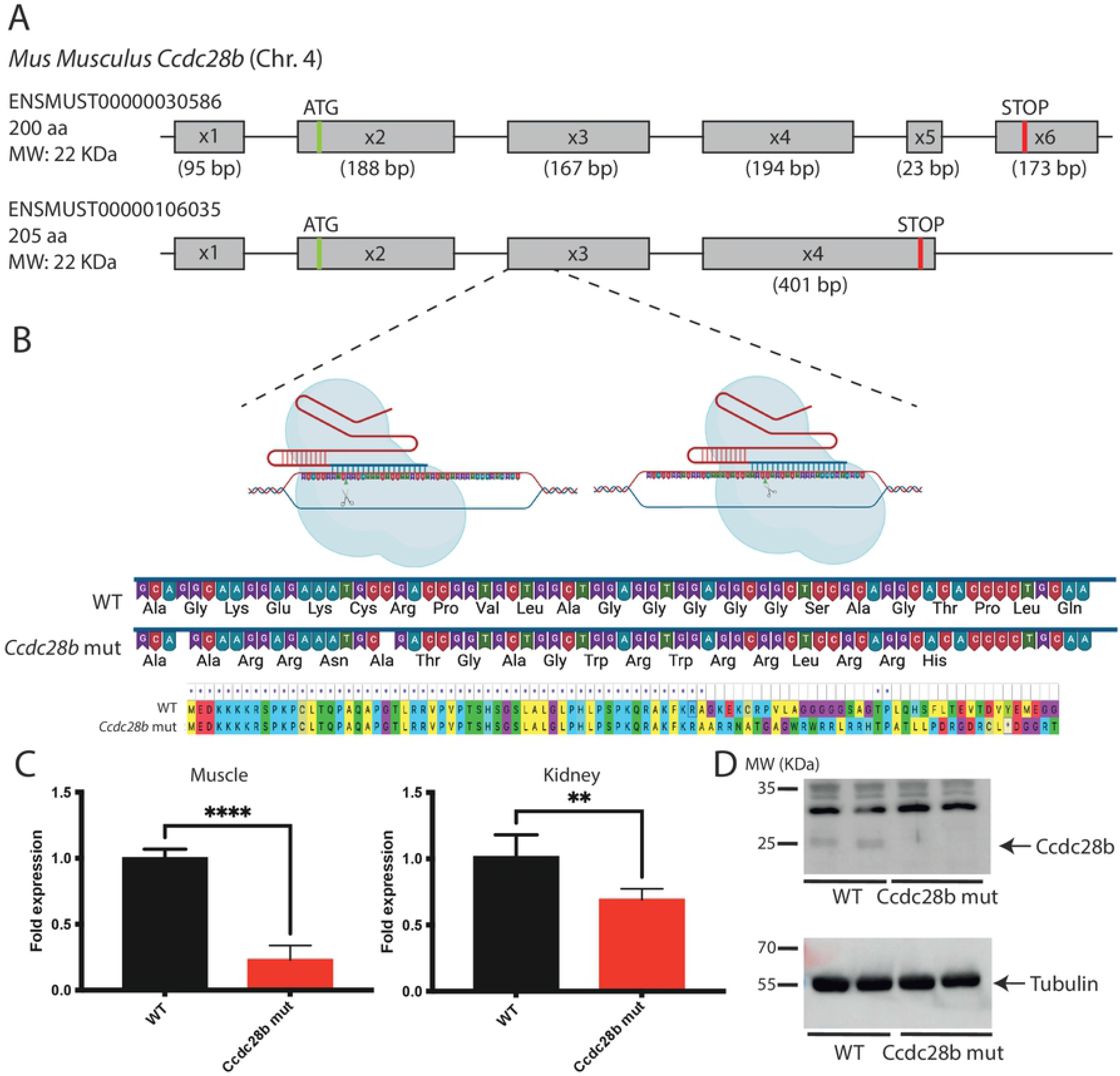
Targeting Ccdc28b in the mouse. **A)** Schematic representation of murine *Ccdc28b* genomic structure, showing exon distribution and the two main reported protein coding ORFs that encode two isoforms differing in their C-terminal sequences. **B)** Graphic representation showing the two gRNAs used in this study targeting the 5’ end of exon 3. The selected mutation comprises two single base deletions leading to a frameshift and the introduction of a PTC. **C)** Real-time quantitative PCR was used to show a significant reduction in *Ccdc28b* mRNA levels, likely due to NMD-mediated degradation of PTC containing mRNA. Expression of *Gapdh* was used for normalization. Results are shown as fold change compared to wt animals. **= *P* < 0.01 and ***= *P* < 0.001. **D)** Western blot analysis showing that a band corresponding to the Ccdc28b expected molecular weight (aprox. 22 KDa) is depleted in tissues of *Ccdc28b mut* animals. Full gels are shown in S2 Fig.

As a first step we aimed to confirm that the expression of *Ccdc28b* was abrogated. As mentioned, the mutation introduces a PTC likely targeting the mRNA for NMD. We analyzed *Ccdc28b* levels by qRT-PCR using specific primers and cDNA obtained from different tissues (*Ccdc28b* is widely expressed; [48]). As expected, we observed a significant reduction in *Ccdc28b* mRNA levels (Fig. 1C). We also PCR-amplified *Ccdc28b* from cDNA of E14 embryos (both *Ccdc28b mut* and C57BL/6J wt) with primers located at the far most 5’ and 3’ ends of the reported isoforms and sequenced the PCR products using Oxford Nanopore technology taking advantage of its long-read sequencing. We detected the expected mutations and did not find any evidence of novel alternative transcripts in *Ccdc28b mut* samples (S1 Fig). Next, we evaluated Ccdc28b at the protein level by western blot: the expected ∼22 KDa Ccdc28b band was absent in the *Ccdc28b mut* samples (Fig 1D; S2 Fig for full gels and muscle blot). Thus, our results indicate that we were able to generate a mutant *Ccdc28b* mouse line (*Ccdc28b mut*).

### Depletion of Ccdc28b does not result in overt ciliary defects but may modulate cilia length in a tissue dependent manner

To begin the characterization of the mutant line we first focus on the known function of Ccdc28b. As previously mentioned, our work in hTERT-RPE cells and zebrafish uncovered a role for Ccdc28b in cilia whereby its depletion resulted in a reduction in ciliary length as well as a reduction in the percentage of ciliated cells [38–40]. We first focused on measuring cilia length and quantifying the proportion of cilia in mouse embryonic fibroblasts (MEFs) obtained from both *Ccdc28b mut* and C57BL/6J (wt) E14 embryos. We quantified the proportion of ciliated cells by counting nuclei and cilia per field, using anti γ-tubulin and anti-acetylated tubulin antibodies to visualize the basal body and ciliary axoneme respectively. We measured cilia length by analyzing at least eight randomly selected fields from each of *Ccdc28b mut* and wt MEFs. Our data showed that depletion of Ccdc28b did not result in shortened cilia nor affected ciliation in these cells: the median cilia length of wt MEFs was 2.33 ± 0.70 μm compared to 2.31 ± 0.78 μm in *Ccdc28b mut* cells (*P* = 0.6522; Fig. 2A-B).

**Figure 2:**
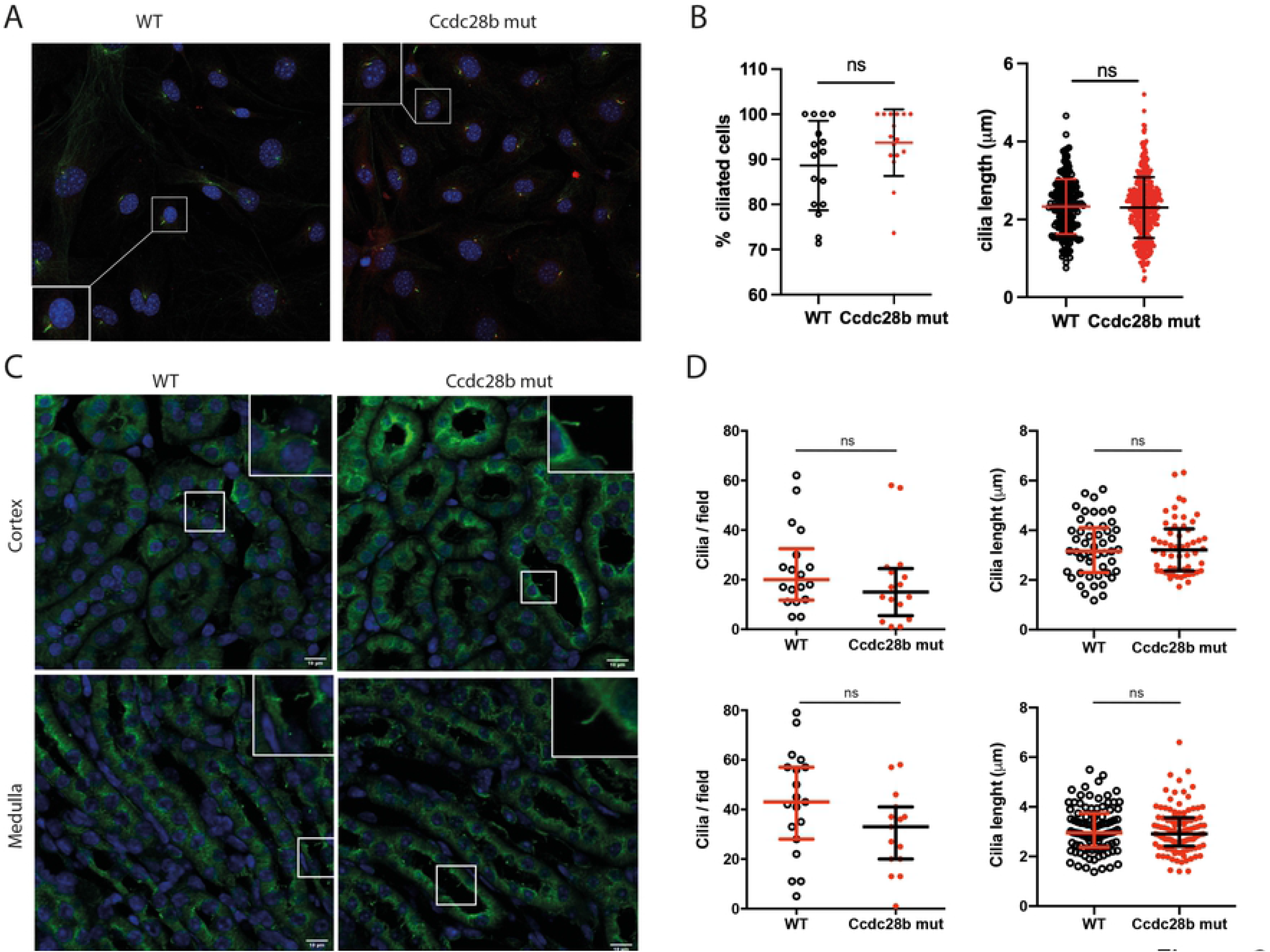
Characterization of cilia in Ccdc28b mut cells and tissue. **A)** Confocal images of wt and *Ccdc28b mut* Mouse Embryonic Fibroblasts (MEFs). DAPI (blue), acetylated tubulin (green) and γ-tubulin (red) were used to visualize nuclei, ciliary axoneme and basal body, respectively. **B)** Cilia length was measured in more than 190 cilia from wt and *Ccdc28b mut* MEFs. Data are presented as individual values plus mean ± SD. **C)** Confocal images of wt and *Ccdc28b mut* kidneys. The upper panels correspond to kidney cortex, and the lower panels correspond to kidney medulla. DAPI (blue) and anti-acetylated tubulin (green) were used to visualize nuclei and cilia respectively. **D)** Quantification of number of cilia per field and cilia length. Data did not have normal distribution and are shown as individual values plus median with interquartile range. Despite no significant differences were found there is a trend towards a reduction in cilia density in both regions in Ccdc28b mut kidneys. ns: not significant.

*Ccdc28b mut* mice reproduce normally although when measuring weeks at first mating *Ccdc28b mut* females showed a delay (approximately 14 weeks) compared to C57BL/6J females (approximately 7 weeks). We then evaluated cilia in the kidney, a tissue where cilia are readily observed. Kidneys from both control and *Ccdc28b mut* animals of 36 weeks of age were processed for immunofluorescence as described in the methods section. No major anatomical or histological differences were observed between *Ccdc28b mut* and wt animals and cilia were readily observed projecting into the lumen of tubules in both genotypes. Again, we quantified the number of cilia per field (assessing areas of comparable cell density) and cilia length. Although we could not find statistically significant differences neither in cilia number or length, we did observe a trend towards a reduction in cilia density in *Ccdc28b mut* preparations, both in the kidney cortex and medulla, compared to wt mice (Fig 2C-D). We also analyzed cilia in the brain. We evaluated amygdala, hippocampus CA1, dentate gyrus (S3 Fig) and performed an in-depth quantification of both cilia density and length in the striatum (Fig 3A-B). Cilia density was comparable between *Ccdc28b mut* and wt controls (Fig 3B). Unexpectedly however, we observed a mild, albeit statistically significant, difference in cilia length whereby cilia in *Ccdc28b mut* animals were longer than controls: 10.16 ± 2.67 μm and 9.38 ± 2.40 μm respectively (*P* = 0.0022; Fig. 3B). Thus, our results show that depletion of *Ccdc28b* does not result in a global cilia defect but can provoke cilia length changes at least in some tissues or cell types.

**Figure 3:**
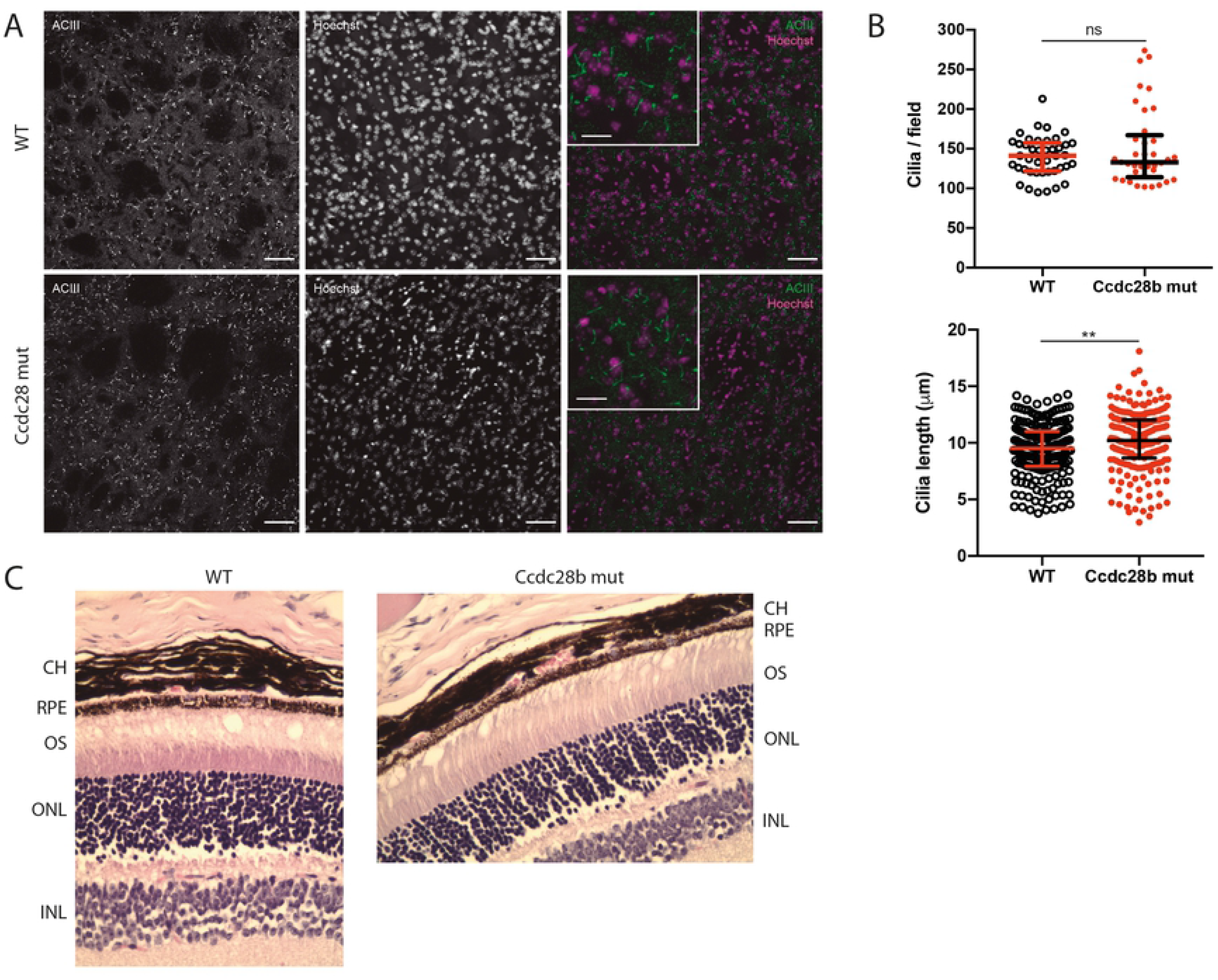
Analysis of brain cilia and the retina in *Ccdc28b mut* animals. **A)** Confocal images showing cilia in the brain striatum. DAPI (magenta) and anti-Type III adenylyl cyclase (ACIII, green) were used to visualized nuclei and ciliary axoneme respectively. Bars in main pictures and insets correspond to 50 and 20 microns respectively. **B)** Quantification of number of cilia per field (at least three pictures per animal, and three animals per genotype) and cilia length (one picture per animal, three animals per genotype) in striatum. Results are represented as individual values plus median with interquartile range. **= *P* < 0.005. **C)** Hematoxylin and eosin stained retinal sections of wt and *Ccdc28b mut* mice. No structural differences were observed in the photoreceptor layer. INL= inner nuclear layer; ONL= outer nuclear layer; OS= Outer segment; RPE= retinal pigment epithelium; CH= choroid.

### Ccdc28b mut animals do not develop retinal degeneration or obesity

To continue the characterization of *Ccdc28b mut* mice we focused on assessing two phenotypes that have been shown to be highly penetrant in both BBS patients [1] and different Bbs mouse models (for example see [44, 46, 49, 50]): retinal degeneration and obesity. In different Bbs mouse models (*Bbs2*, *Bbs4* and *Bbs12*) retinal degeneration has been shown to be progressive, first evident as a thinning of the outer nuclear layer (photoreceptors) by as early as 6 weeks of age and characterized by complete loss of the outer segment in older animals (7 months old in Bbs4; [49]). In contrast, *Ccdc28b mut* retinas in both 12 weeks and 9 months old animals presented normal structure including the photoreceptor layer (Fig 3C; 9-month-old retinas are shown).

*Bbs2*, *Bbs4*, *Bbs6*, and *Bbs12* KOs, as well as a *Bbs1^M390R/M390R^* knock-in animals, develop obesity driven by hyperphagia [44-46, 49-51]. To assess whether our *Ccdc28b mut* animals present similar phenotypes we measured i) weight gain on normal diet, ii) weight gain on high fat diet (HFD), iii) food consumption and iv) systemic glucose handling (Fig 4). Bbs mutant animals have been reported to be runted at birth and then rapidly start gaining weight at an increased rate. *Ccdc28b mut* animals presented a normal appearance at birth and gained weight at comparable rates to control animals when fed *ad libitum* on a normal diet. Mice were followed up to 21 weeks of age (Fig 4A). Next, we generated two groups of animals (n= 7 per genotype) and fed them a HFD starting at 8 weeks of age. We did not observe significant weight differences between *Ccdc28b muts* and wt animals (Fig 4B-C). Accordingly, studying our mice in metabolic cages did not reveal any signs of increased food intake (Fig 4D-E). We also evaluated systemic glucose management by measuring glucose blood levels (basal glycemia) after 16 hours of starvation and performing glucose tolerance tests (GTT) before starting the HFD and at two time points during the treatment (7 and 11 weeks in HFD respectively). *Ccdc28b mut* mice showed significantly elevated basal glucose levels after 7 weeks on HFD (Fig 4F). By 11 weeks of HFD the difference was lost but mainly due to an increase in the glucose basal levels in control animals (Fig 4F and S4 Fig). In the GTTs we did not observe statistically significant differences although *Ccdc28b mut* mice showed a trend towards an impaired GTT response, particularly after 11 weeks on HFD (Fig 4G-I). Overall, our results show that *Ccdc28b mut* animals do not present hyperphagia or obesity but do show a mild phenotype related to systemic glucose management.

**Figure 4:**
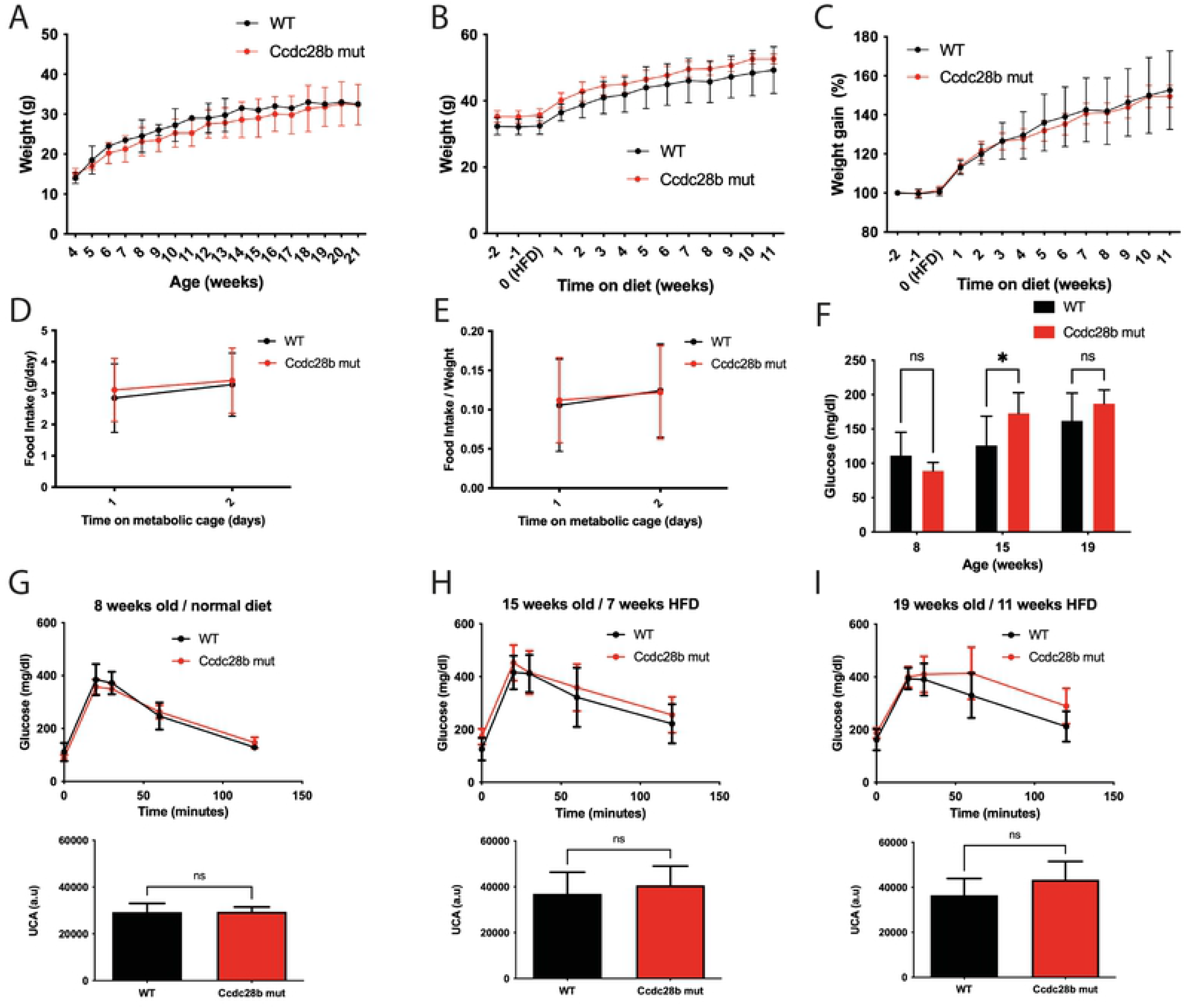
Metabolic characterization of *Ccdc28b mut* mice. **A)** Growth curve of *Ccdc28b mut* and wt littermates during a ND *ad libitum*. **B)** Growth curve of *Ccdc28b mut* and wt littermates during a HFD *ad libitum*. Animals at week -2 of diet were 6 weeks old. **C)** Weight gain curve of animals shown in B. Weight is normalized to the value at week -2. **D)** Food intake measured in metabolic cages. **E)** Food intake showed in D normalized to body weight. In all sections error bars correspond to mean and SD. **F)** Basal glycemia measured in the corresponding GTT tests (panels G-I) measured at 8, 15 and 19 weeks of age. *= *P* < 0.05. **G-I)** GTT results for *Ccdc28b mut* and wt controls at 8 weeks of age and normal diet (G), at 15 weeks of age with 7 weeks of HFD (H) and at 19 weeks of age after 11 weeks of HFD (I). Both glucose level curves and the quantification of the area under the curve are shown as mean ± SD of the analyzed animals.

### Ccdc28b KO animals present autism-like behavioral phenotypes

Behavioral phenotypes have been reported in BBS patients [1, 52]. In agreement with this documented observations, social dominance defects and anxiety related responses have been well documented in different Bbs mouse models using standardized tests such as open field, light-dark box test and social dominance tube test [44–47]. In contrast, compulsive obsessive behavior and phenotypes related to autistic spectrum disorder have also been observed in BBS patients [1, 52] but have been less studied in mouse models. Importantly, Kerr and colleagues assessed the behavioral phenotypes in a cohort of twenty-four confirmed BBS patients demonstrating a high incidence of symptoms associated with autism [42]. In this context, we asked whether *Ccdc28b mut* mice displayed behaviors that could be relevant to BBS.

We started our analysis of *Ccdc28b mut* animals performing an open field test to assess exploratory activity and overall movement. We analyzed both female and male animals and did not find significant differences in total distance traveled between *Ccdc28b mut* and wt mice (Fig 5A). We also evaluated time freezing and time in the periphery versus center of the field where we did not find significant differences between genotypes on either females or males (Fig 5B-C). Next, we evaluated anxiety directly by performing the elevated plus maze test (EPM). *Ccdc28b mut* animals spent comparable amounts of time, and traveled comparable distances, in the open arms as control animals (Fig 5D). To directly test for alterations in hippocampal functions we performed a Novel Object Recognition (NOR) test. In this assay we evaluated memory by presenting individual animals with two objects for 10 min, and 24 hours later exchanging one known object for a novel one: the number of interactions of each mouse with each object (known *vs* novel) was quantified as described in methods. Both *Ccdc28b mut* and wt controls showed a significantly higher number of interactions with the novel object (Fig 5E).

**Figure 5:**
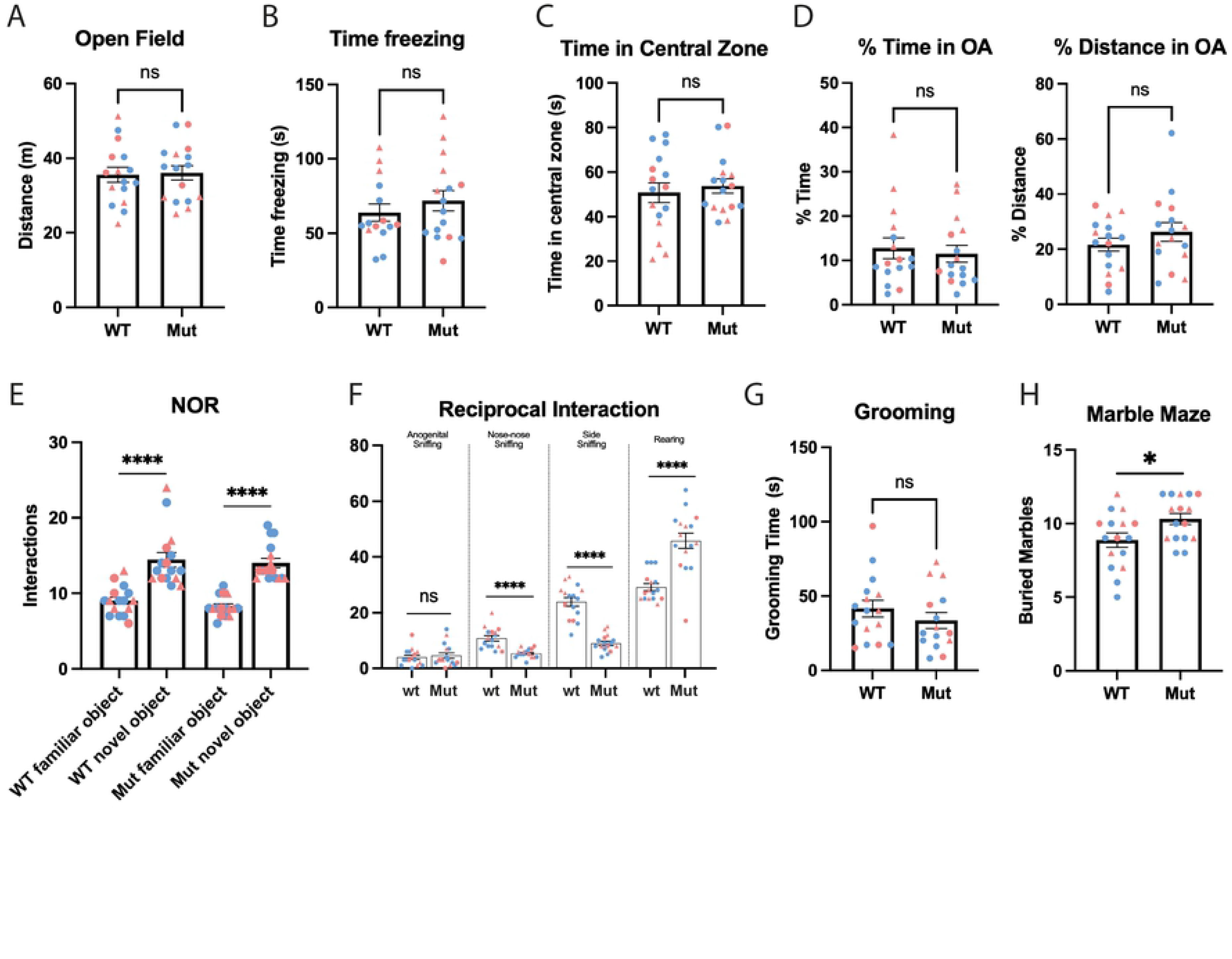
Evaluation of behavioral and social phenotypes in *Ccdc28b mut* mice. An open-field test was performed, and different parameters were scored: total distance traveled (animals of both sexes are shown together with different colors) **(A)**, time freezing **(B)** and time in the periphery *vs* center of the field **(C).** No differences were observed between wt and *Ccdc28b mut* mice. **D)** Results from elevated plus maze test showed no signs of anxiety as wt and *Ccdc28b mut* mice spent comparable amounts of time, and travelled comparable distances, in the open arms. **E)** The Novel Object Recognition test showed that there is no alteration in memory in *Ccdc28b mut* mice, as they presented an equally higher number of interactions with novel objects when compared to wt mice. **F)** Reciprocal Social Interaction Test resulted in significant differences in nose-nose, side sniffing, and rearing, consistent with an Autism Spectrum Disorder (ASD) related phenotype, while no differences were found in anogenital sniffing. **G)** No differences were observed in grooming time between *Ccdc28b* and wt animals. **H)** Marble burying test showing that *Ccdc28b mut* buried more marbles than wt animals, a behavior consistent with an obsessive-compulsive phenotype. In all graphs where female and male mice are shown together, light blue indicates male and light red shows females. Triangles were used to indicate the five females analyzed in the second cohort (see Methods). All error bars correspond to mean and SEM. *= *P* < 0.05; ****= *P* < 0.0001.

Next, to study the impact of *Ccdc28b* in Autism Spectrum Disorder (ASD) relevant behaviors, we performed a reciprocal social interaction test, a social dyadic test where the interrogated animal is presented with a previously unknown mouse to then quantify natural social behaviors. In this assay we evaluated anogenital, nose-nose and side sniffing, as well as grooming and rearing, measuring the number (frequency) of such behaviors in a period of 10 minutes. While *Ccdc28b mut* and wt did not differ in the frequency of anogenital sniffing or grooming, significant differences were readily observed for the other parameters: *Ccdc28b mut* animals presented a significant reduction in nose-nose and side sniffing and, in agreement with those results, also increased rearing (Fig 5F). Finally, we assessed stereotypical behaviors. Whereas *Ccdc28b mut* animals did not show differences in grooming when compared to controls (Fig 5G), *Ccdc28b mut* animals consistently buried more marbles than control animals in the marble burying test, a phenotype that was clearly observed both in males and females (Fig 5H). Altogether, our results show that while *Ccdc28b mut* mice neither present defects related to movement, grooming, nor signs of memory loss, they do show obsessive compulsive and ASD-related phenotypes.

### CCDC28B mutations in individuals with autism

We next examined the effect of deleterious mutations in *CCDC28B* among children with autism. We analyzed copy-number variation and single nucleotide variants (SNVs) involving *CCDC28B* among 2,532 individuals with autism and their families from the Simons Simplex Collection (SSC). We identified seven probands with rare, deleterious SNVs in *CCDC28B* that were present at <0.001 frequency in the gnomAD control database, and one proband carried a duplication encompassing *CCDC28B* (Table 1). We also analyzed quantitative phenotypes of *CCDC28B* mutation carriers (Table 2) and compared them to a distribution of average scores derived from 1000 random sampling of equivalent numbers of probands from the cohort (see Methods). While quantitative measures of autism phenotypes such as intelligence quotient, child and adult behavioral meassures, and repetitive behavior among these probands were not different from the rest of the affected individuals from the SSC cohort, *CCDC28B* mutation carriers showed a significantly reduced BMI (empirical *P*= 0.04) than expected for the entire cohort and a milder social responsiveness raw score (empirical *P*= 0.04) (S5 Fig). These results suggest that mutations in *CCDC28B* lead to milder autism features.

**Table 1:** *CCDC28B* mutations present in probands in the SSC.

| **Sample** | **Chr** | **Position** | **Ref** | **Alt** | **Mutation Type** | **CADD** | **gnomAD Frequency** | **Inheritance** |
| --- | --- | --- | --- | --- | --- | --- | --- | --- |
| **Proband 1** | **chr1** | **32201979** | **C** | **T** | **missense** | **22.9** | **0.00002** | **Father** |
| **Proband 2** | **chr1** | **32201988** | **C** | **T** | **missense** | **22.9** | **0.000056** | **Mother** |
| **Proband 3** | **chr1** | **32204030** | **C** | **T** | **missense** | **33** | **0.000015** | **Mother** |
| **Proband 4** | **chr1** | **32204245** | **C** | **T** | **missense** | **33** | **0.00002** | **Father** |
| **Proband 5** | **chr1** | **32204360** | **T** | **C** | **missense** | **26.9** | **N/A** | **De novo** |
| **Proband 6** | **chr1** | **32204362** | **G** | **A** | **missense** | **28.3** | **N/A** | **De novo** |
| **Proband 7** | **chr1** | **32204601** | **G** | **A** | **missense** | **24** | **0.000005** | **Mother** |
| **Proband 8** | **chr1** | **31125281 - 3607897** | **-** | **-** | **duplication** | **N/A** | **N/A** | **Mother** |

**Table 2.**
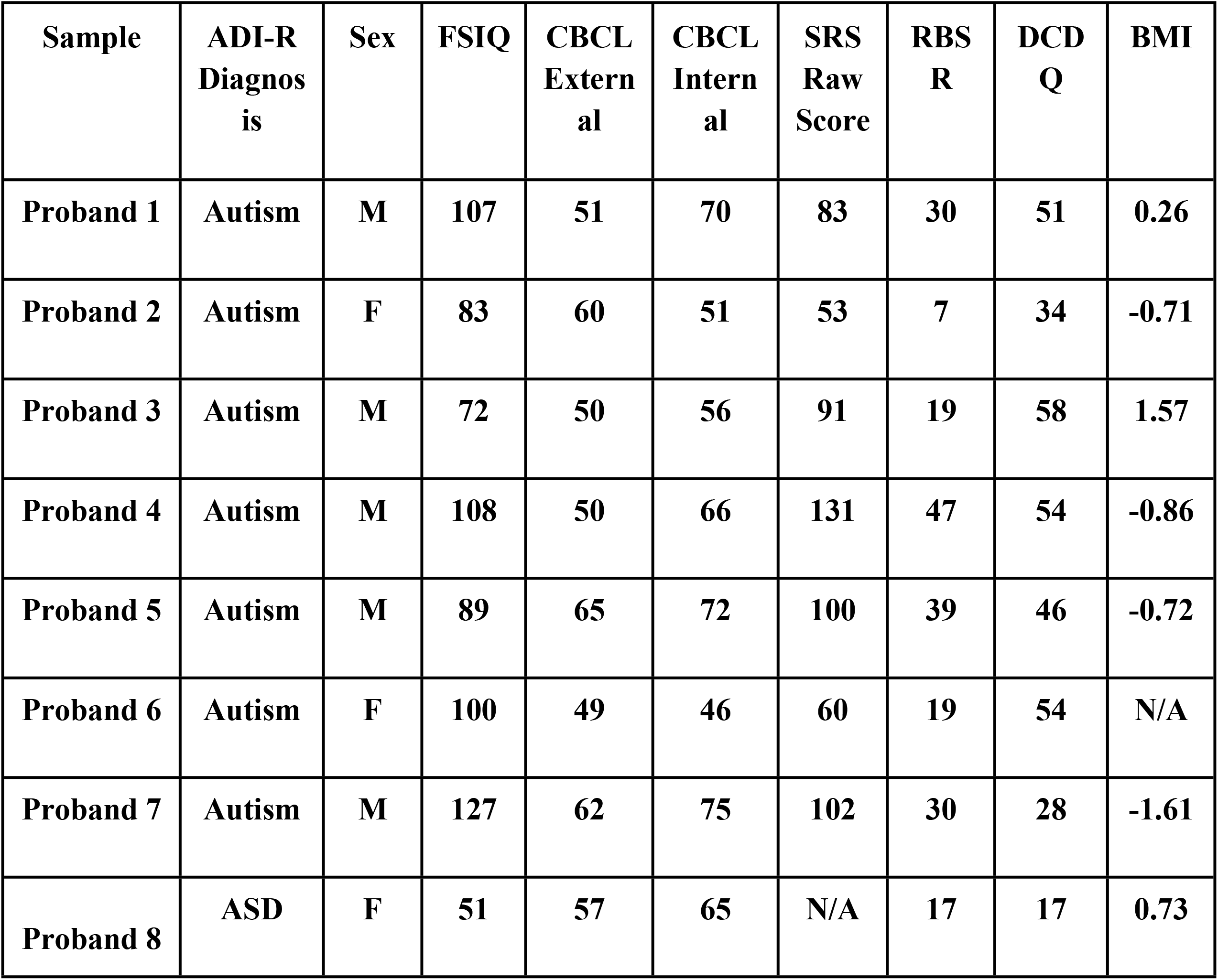
Phenotypes in SSC probands carrying *CCDC28B* mutations.

| **Sample** | **ADI-R**<br>**Diagnosis** | **Sex** | **FSIQ** | **CBCL**<br>**External** | **CBCL**<br>**Internal** | **SRS**<br>**Raw**<br>**Score** | **RBS**<br>**R** | **DCD**<br>**Q** | **BMI** |
| --- | --- | --- | --- | --- | --- | --- | --- | --- | --- |
| **Proband 1** | **Autism** | **M** | **107** | **51** | **70** | **83** | **30** | **51** | **0.26** |
| **Proband 2** | **Autism** | **F** | **83** | **60** | **51** | **53** | **7** | **34** | **-0.71** |
| **Proband 3** | **Autism** | **M** | **72** | **50** | **56** | **91** | **19** | **58** | **1.57** |
| **Proband 4** | **Autism** | **M** | **108** | **50** | **66** | **131** | **47** | **54** | **-0.86** |
| **Proband 5** | **Autism** | **M** | **89** | **65** | **72** | **100** | **39** | **46** | **-0.72** |
| **Proband 6** | **Autism** | **F** | **100** | **49** | **46** | **60** | **19** | **54** | **N/A** |
| **Proband 7** | **Autism** | **M** | **127** | **62** | **75** | **102** | **30** | **28** | **-1.61** |
| **Proband 8** | **ASD** | **F** | **51** | **57** | **65** | **N/A** | **17** | **17** | **0.73** |

## DISCUSSION

*CCDC28B* was first identified as a second site modifier of BBS whereby a reduction in its levels, in conjunction with mutations in *bona fide* BBS genes, was shown to result in a more severe presentation of the syndrome in some families [35]. Also, we have shown previously that *CCDC28B* participates in the regulation of cilia in both cells and *in vivo* in zebrafish: depletion of *ccdc28b* resulted in cilia shortening and defective ciliogenesis in different zebrafish tissues and organs [38–40]. Consequently, knockdown of *ccdc28b* in the fish using primarily a morpholino approach resulted in several phenotypes that are characteristic of Bbs and other ciliary mutants, such as a curved body axis, pigmentation defects, craniofacial malformations, and hydrocephaly ([38, 39] and references within). Therefore, while our previous results provided important functional information to understand CCDC28B biological role, they also led us to hypothesize that null mutations in *CCDC28B* could be sufficient to cause a ciliopathy such as BBS, or even the more severe Meckel-Gruber syndrome (MKS), two conditions with a high degree of genetic overlap [22, 38]. Importantly, the mutation described in patients, and shown to modify the presentation of the syndrome, is likely a hypomorphic mutation caused by a synonymous change affecting *CCDC28B* mRNA splicing thus resulting in reduced, but not abolished, CCDC28B levels [35].

We therefore decided to target *Ccdc28b* in the mouse and performed a characterization directing our attention to different phenotypes that have been reported previously for this gene (effect on cilia) and for BBS models. Overall, our data indicate that targeting *Ccdc28b* in the mouse is not sufficient to cause a strong ciliary defect and accordingly, does not cause phenotypes that are highly penetrant in different mouse Bbs models, such as retinal degeneration or the development of obesity. We cannot rule out at this point the possibility of *Ccdc28b mut* animals presenting a predisposition to develop BBS like phenotypes. Older animals (we assessed mice up to 9 month of age) will have to be evaluated to test this possibility.

Our results are fully compatible with a second site modifier role for *CCDC28B*. *Ccdc28b mut* mice did not show signs of retinopathy and did not develop obesity at least up to nine months of age. Moreover, using metabolic cages we were able to show that *Ccdc28b muts* are not hyperphagic. Interestingly however, *Ccdc28b mut* mice show both a trend towards presenting a worst performance than controls in the GTTs and presented significantly higher glucose basal levels after 7 weeks on HFD. Thus, our results suggest a mild phenotype related to glucose handling. Interestingly, the consequences of BBS gene mutations on systemic glucose management are still not entirely clear. For example, while *Bbs4* KO animals have been shown to present impaired glucose handling, mainly due to defective insulin secretion [59], *Bbs12* KO animals have been shown to present an improved glucose metabolism [50]. Likewise, reports are showing that some BBS patients could present a dissociation between obesity and the development of type II diabetes [50, 60]. Thus, it will be interesting to study whether variants in genes such as *CCDC28B* could contribute to modulating the presentation of obesity in BBS. Crossing this new *Ccdc28b mut* mouse line with available Bbs mutants will allow to start tackling this issue.

This work also underscores differences with our previous data working on cells and zebrafish. Besides differences in models (culture cells *vs* zebrafish *vs* mouse), one possibility is that our *Ccdc28b mut* animals are not complete functional knockouts. We believe this to be unlikely considering our qRT-PCR, sequencing, and western blot results. Also, the mutant *Ccdc28b* ARNm could encode an 88 amino acid polypeptide which would only include the first 56 amino acids of Ccdc28b. We favor a second scenario which relies on genetic compensation, a mechanism that has been shown to be particularly important to understand differences between targeting genes at the mRNA level versus at the genomic level [53, 54]. This phenomenon has been well documented in zebrafish and other model organisms where the same gene has been targeted, for example, using morpholinos to block mRNA splicing/translation, and through genome editing, highlighting the complexity and plasticity of the genome. Oftentimes the phenotype of morphants is significantly more severe than that of genomic mutants and different mechanisms have been shown to explain these differences. For example, CRISPR-Cas9 mutant lines have been shown to present altered mRNA splicing thus bypassing the mutated residue/exon [55]. As mentioned, we could not find any evidence for alternative splicing in our mice. Importantly however, the compensation can also occur by rescuing the cellular function rather than a particular gene. For example, *egfl7* (endothelial extracellular matrix gene) genomic mutants, but not morphants, upregulate genes that are functionally related to *egfl7* [54]. Similarly, while targeting *CEP290* (*NPHP6*), a ciliopathy gene linked to BBS, in zebrafish at the mRNA level resulted in severe cilia-related phenotypes, only a mild defect restricted to photoreceptors was observed in genomic mutants. Interestingly, the authors found that the mild presentation of the later was associated with the upregulation of several genes associated to ciliary function [56]. Importantly, it has been shown that upregulation of compensatory genes is triggered by mutations that introduce a PTC in a mechanism that relies on NMD [57, 58]. Therefore, rather than a weakness of the *Ccdc28b mut* mouse model, its mild phenotype could be seen as an opportunity to perform transcriptomic/proteomic studies attempting to dissect this compensation mechanism. Such data will likely shed important insight to understand the biological role of *CCDC28B* and the cellular/molecular pathways involved in cilia regulation.

Finally, we did observe a clear behavioral phenotype in our *Ccdc28b mut* animals. Phenotypes such as anxiety, social dominance and associative learning defects, have been reported in different BBS mouse models [44–47]. For example, it was shown that BBS mice present problems in context fear conditioning due to impaired neurogenesis [47]. Our *Ccdc28b mut* animals did not show differences with controls in the open field, the EPM, or the NOR tests, thus ruling out significant locomotor, anxiety, and memory dysfunctions. *Ccdc28b mut* animals however did show clear social interaction defects as well as stereotypical phenotypes suggestive of an ASD-like behavior. Interestingly, ASD-like behaviors have been described in BBS patients (see for example [41–43]). Moreover, although historically reported as a rare presentation, a comprehensive study of behavioral phenotypes in twenty-four BBS patients even determined that autism related symptoms were present in 77% of cases [42]. Thus, it is tempting to speculate that decreased CCDC28B function could contribute to modulate the penetrance, the expressivity, or both, of ASD-like phenotypes in BBS patients, a possibility that will require further studies.

Interestingly, our results also suggest that *CCDC28B* could contribute causal alleles in ASD patients in a BBS-independent context: filtering for rare, likely deleterious variants, we identified eight SSC probands carrying *CCDC28B* mutations. Furthermore, by analyzing the phenotype of these probands in comparison with a randomly sampled group of SSC probands, our results suggest that mutations in *CCDC28B* could lead to milder forms of autism. Thus, this *Ccdc28b mut* mouse model could provide an opportunity to study cellular/molecular aspects of ASD phenotypes. In our analysis of the brain striatum, we observed that cilia were longer than controls. While this elongation was subtle, the difference was statistically significant. Two points regarding this cilia phenotype. First, all our previous data have clearly shown that Ccdc28b plays a pro-ciliogenic role, whereby its depletion results in shortened cilia. Thus, one possibility is that the observation of longer cilia in striatum could be underscoring differences between cell types. It is tempting to speculate however that this slightly elongated cilia could be a consequence of genetic compensation and the upregulation of pro-ciliogenic genes in the absence of Ccdc28b. Further studies will be required to test this intriguing possibility. The second point to discuss here relates to the potential physiological relevance of this finding. Wang and colleagues recently published a work using induced pluripotent stem cell-derived (iPSC) neurons obtained from both BBS patients and controls where they show that mutations in BBS genes affect neurite outgrowth and neuronal energy homeostasis. Interestingly, BBS mutant iPSC-derived neurons presented elongated cilia [61]. Observing elongated cilia in a region of the brain that has been linked to the development of ASD conditions, like the striatum [62, 63], suggests the possibility of cilia playing an active role in the development of these conditions. Further work will be needed to determine whether this is the case, and our *Ccdc28b mut* model will likely be an important reagent in this context. More broadly, the mouse line presented here, together with other already available models, will likely contribute to further our still incomplete understanding of cilia in neuronal homeostasis, brain development, the establishment of neuronal circuitry and its impact on behavior [64–66]. Further dissecting the biological role of CCDC28B will likely shed important insight to understand not only BBS but also ASD.

## METHODS

### Animals

All animal procedures to generate the mutant line were performed at the SPF animal facility of the Laboratory Animal Biotechnology Unit of Institut Pasteur de Montevideo. Experimental protocols were opportunely approved by the Institutional Animal Ethics Committee (protocol number 007-18), in accordance with national law 18.611 and international animal care guidelines (Guide for the Care and Use of Laboratory Animal; [67]) regarding laboratory animal’s protocols. Mice were housed on individually ventilated cages (Tecniplast, Milan, Italy) containing chip bedding (Toplit 6, SAFE, Augy, France), in a controlled environment at 20 ± 1°C with a relative humidity of 40-60%, in a 14/10 h light-dark cycle. Autoclaved food (Labdiet 5K67, PMI Nutrition, IN, US) and autoclaved filtered water were administered *ad libitum*.

Cytoplasmic microinjection was performed in C57BL/6J zygotes using a mix of 30 ng/µl Cas9 mRNA (Synthego, Menlo Park, CA, US), and 15 ng/µl of each sgRNA (2 guides were used) (Synthego), diluted in standard microinjection buffer. Viable embryos were transferred into the oviduct of B6D2F1 0.5 days post coitum (dpc) pseudo-pregnant females (25 embryos/female in average), following surgical procedures established in our animal facility [68]. For surgery, recipient females were anesthetized with a mixture of ketamine (100 mg/kg, Pharmaservice, Ripoll Vet, Montevideo, Uruguay) and xylazine (10 mg/kg, Seton 2%; Calier, Montevideo, Uruguay). Tolfenamic acid was administered subcutaneously (1 mg/kg, Tolfedine, Vetoquinol, Madrid, Spain) to provide analgesia and anti-inflammatory effects [69]. Pregnancy diagnosis was determined by visual inspection by an experienced animal caretaker two weeks after embryo transfer, and litter size was recorded on day 7 after birth. Pups were tail-biopsied and genotyped 21 days after birth, and mutant animals were maintained as founders.

### RNA Isolation and qRT-PCR

Tissues were homogenized in TRIzol (Invitrogen) for RNA extraction according to the manufacturer’s protocol. Reverse transcription was done using SuperScript II RT (Invitrogen) and qRT-PCR was performed using SYBR FAST Universal 2X qPCR Master Mix (Kapa) on a QuantStudio 3 RT-PCR System (Thermo Fisher Scientific). All samples were run in triplicate and the CT value was normalized to calculate relative expression of each gene. The fold expression was calculated using the ΔΔCt method with *Gapdh* as a reference gene.

### Western Blotting

Tissues were lysed using RIPA buffer (25mM Tris pH 8.0, 150mM NaCl, 1% NP-40, 0.1% SDS, 1% sodium deoxycholate) supplemented with a protease inhibitor cocktail (Sigma). Protein concentrations were determined using the BCA Protein Assay Kit (Thermo Fisher Scientific) and 100µg of total protein were loaded into SDS-PAGE gels, transferred to PVDF membranes and probed with anti-CCDC28B (Invitrogen, 1/1000) overnight at 4°C. HRP-conjugated secondary antibody was used.

### Cell culture and immunofluorescence

Mouse embryonic fibroblasts (MEFs) were obtained from E13.5-E14.5 following standard procedures and maintained sub-confluent in DMEM Glutamax with 10% FBS, Hepes 10mM, penicillin 10000 U/mL and streptomycin 10000 μg/mL(Maintenance Medium, MM) under controlled conditions at 37° C with 5% CO_2_. For immunofluorescence studies cells were cultured on glass coverslips and at 80% confluency, MM was replaced with medium containing 0.5% FBS for 24 hr to stimulate ciliation. Cells were fixed with 4% paraformaldehyde (PFA) in 0.1 M phosphate buffer saline (PBS), permeabilized with 0.1% Triton-X100, blocked with 5.5% FBS and stained with anti-gamma and anti-acetylated tubulin primary antibodies (Sigma) followed by the corresponding secondary antibodies conjugated to AF488 or TMRM (Invitrogen). Nuclei were stained with DAPI (Invitrogen). Images were taken in a Zeiss LSM 880 confocal microscopy. Eight randomly selected confocal fields from cultured MEFs from at least two *Ccdc28b mut* and two wt embryos were analyzed. Cilia length was measured using the freehand ROI selection tool of the FIJI image processing package [70]. The proportion of ciliated cells was calculated by counting the number of cells showing γ-tubulin and anti-acetylated tubulin staining over the total number of nuclei.

### Brain, kidney and eye histological and immunofluorescence analysis

The perfusion of the mice was performed during the light (resting) phase of the sleep-wake cycle (between 12:00 and 16:00 h, lights on at 7:00). The animals were anaesthetized with ketamine/xylacine (90 and 14 mg/kg, respectively) and perfused with PBS followed by 4% PFA. Brains, kidneys and eyes were immediately dissected out and fixed by immersion in 4% PFA overnight (ON). Thereafter, the brains were cryoprotected in 30% sucrose solution in 0.1 M PBS for 48 h and frozen. Coronal sections (30 μm) were obtained by a cryostat (Leica CM 1900, Leica Microsystems, Nussloch, Germany). Sections containing, hippocampus, amygdala and striatum (based on The mouse brain atlas; Paxinos & Franklin 2000) were collected and stored in an anti-freeze solution at −20 °C until immunostaining procedures were performed. Kidneys and eyes were embedded in paraffin and cut into 6 micron slices. Eye sections were dewaxed, rehydrated and stained with Hematoxilin and Eosin following standard procedures.

Cilia were detected in brain and kidney sections by immunofluorescence. Renal cilia were stained as previously described [71], using boiling in 10 mM citrate buffer, pH6, for antigen retrieval and anti-acetylated tubulin (Sigma) 1/300 in PBS containing 0.5% of normal goat serum and 0.05% Tween20, for cilia detection. Goat anti-mouse Ig coupled to AF488 (Thermo) 1/1000 was used as secondary antibody and DAPI (Thermo) 1/5000 for nuclear staining. Cilia staining in brain sections was performed by detecting Type III adenylyl cyclase (AC-III). Briefly, free-floating sections were incubated with rabbit anti-AC-III primary antibodies 1/500 (Santa Cruz Biotechnologies) in PBS plus 0.3% Triton (PBS-T) and normal donkey serum (NDS) 1.5% for 48 h at 4 °C. Then, the sections were incubated with biotinylated donkey anti-rabbit (DAR) 1/600 (Jackson ImmunoResearch) in PBS-T and NDS 3% for 90 min. Afterward, they were incubated with streptavidin-Alexa fluor 555 conjugate 1/2000 (Molecular Probes) in PBS for 2 h. Finally, the sections were mounted in Superfrost Plus slides with Vectashield (Vector Labs) and Hoechst was included to visualize nuclei. Negative controls consisted of omission of the primary antibodies. We obtained three 20x microphotographs of each region of interest. Images were obtained in a confocal microscopy Zeiss LSM 880 using a 63x oil objective.

### *In vivo* metabolic studies

The animals used in this study were raised and maintained according to standard protocols that were approved by the ethical committee at the Institut Pasteur de Montevideo (protocol number 003-19). Mice were fed *ad libitum*, first with a normal control diet (ND, Labdiet 5K67, PMI Nutrition, IN, US) and then with a high-fat diet (HFD, Test Diet 5TJN). Body weight was recorded weekly. For the weight gain control in ND a group of 4 wt (2 males and 2 females), and 7 *Ccdc28b mut* (4 males and 3 females) mice were followed up from week 4 to week 21. For weight gain in HFD a group of 7 wt and 7 *Ccdc28b mut* (all male) were fed with ND until week 8, then changed to HFD until week 21 when all mice were euthanized. For glucose tolerance testing, mice were starved for 16 h before receiving a single intraperitoneal glucose injection (1,5 g/kg). Glycemia was measured from tail vein blood using a hand-held glucometer (Accu-Chek, Roche). For food intake measurements, mice were transferred to metabolic cages, and after 24 h of adaptation, food was weighed every 24 h for three consecutive days.

### Behavioral Analyses

Behavioral assays were conducted in a total of 32 mice, all between 11 and 13 weeks old, divided as follows: 8 wt males, 8 *Ccdc28b mut* males, 8 wt females and 8 *Ccdc28b mut* females. Cohort 1 consisted of all 16 males and 3 females of each group, while cohort 2 consisted of 5 females of each group. All assays were recorded using Any-maze software, except the marble maze test, and for all assays where automatization was not possible, videos were scored by investigators blinded to genotype.

### Open Field

Locomotor activity was assayed in an open field opaque white plexiglass chamber (39×60×50 cm), where animals were left for 6 min sessions to explore freely and afterwards were returned to their home cages. Any-maze software was used to measure total distance, time freezing and time in central area.

### Marble Maze

Marble maze test can be used to assess repetitive, compulsive-like behaviors [72]. Briefly, each individual mouse was placed in an arena (15×27×20 cm) for 30min, with a 5cm layer of clean bedding, where 12 opaque glass marbles were distributed evenly. To be scored as buried, marbles had to be at least 50% covered by bedding.

### Grooming

Grooming can be used as a repetitive behavior assay [73]. Cages were left without bedding to eliminate digging, which can be a competing behavior [74]. Mice were placed individually in standard mouse cages (15×27×13 cm). Sessions lasted 20 min, with the first 10 min being unscored as a habituation period. During the second 10 min of the session, cumulative time spent grooming was scored manually by an investigator uninformed of genotype.

### Reciprocal Interaction

The reciprocal interaction test was used to assess social behaviors and interactions towards an age- and sex matched partner. To this end, mice were placed in standard mouse cages (15×27×13 cm), and interactions were recorded for 10 min, which is the period during which most social interactions happen [74]. The test was recorded using any-maze software and scored manually. Parameters of social behaviors measured were anogenital sniffing, nose-nose sniffing, side sniffing, self-grooming, and rearing [75, 76].

### Elevated Plus Maze

The Elevated Plus-Maze test is used as a model of anxiety [77, 78]. The apparatus consists of four arms (29×7), two of them with walls 16cm high (closed arms), and the other two have no walls, the open arms, the center that connects them is 8×8cm. This maze was placed 50cm from the floor, and mice were placed in the middle, facing an open arm and left to explore freely for 5 min. Time and distance spent in open arms was scored using Any-maze software.

### Novel object recognition

This test is constructed to assess the mouse’s ability to recognize a novel object in the environment, and is divided in three phases: habituation, familiarization, and test phase [79]. Briefly, on the first day, for the habituation period, animals were placed in an empty arena (25×25×35 cm) and left to explore for 5 min before returning to their home cages. 24 h later, the familiarization phase was performed, in which two identical objects were placed in the arena, and each mouse was allowed to explore for 10 min. Finally, on the third day, the test phase consisted of the mouse in the same arena for 10 min, where one of the objects was switched for a novel object, previously unseen to the animal. During the last two phases, objects were placed in opposite and symmetrical corners of the arena [79, 80]. Behavior was scored for the first 2 min of the test phase, or until the mouse had interacted with both objects a total of 20 times.

### Analysis of the Simons Simplex Collection cohort

We assessed for copy-number variants identified from microarrays and single nucleotide variants (SNV), including missense, loss of function, and splice-site mutations affecting *CCDC28B* from whole genome sequencing of 2,532 individuals from the Simmons Simplex Collection (SSC) cohort. We filtered SNV calls to only include variants predicted to be deleterious (with a CADD score >20). We analyzed seven different available quantitative phenotypes, including FSIQ, CBCL External, CBCL Internal, SRS raw score, RBS-R, DCDQ, and BMI z-scores. To test whether the average score observed in the probands with *CCDC28B* mutations was different than expected, we generated 1000 random samples of probands from the SSC that were of the same sample size as the number of probands *CCDC28B* mutations (7 or 8 depending on the phenotype) and calculated the average phenotype score for each of these 1000 simulated samples. We then compared the average of the *CCDC28B* probands with the generated distribution and calculated (a) the number of standard deviations away from the sampled population mean (z scores), and (b) the proportion of samples with a phenotype score at least as extreme as the one observed in the *CCDC28B* mutation probands (empirical p).

### Statistical Analyses

All data were analyzed using GraphPad Prism 9. Unpaired t-test was used to compare the fold expression in the qRT-PCR analysis. For GTT analysis, the area under the curve was calculated in GraphPad Prism 9, and then compared between both groups using the unpaired t-test. Basal glucose between both groups at different times was compared with the 2-way ANOVA with multiple comparisons. The proportion of cilia positive cells was calculated by counting the number of cilia with clear acetylated-tubulin signal over the total number of nuclei and the comparison between the different samples was performed using a test of Hypothesis specific for comparison of two proportions (hypothesis test for proportions). For the analysis of cilia length, and number of cilia per field, we first tested the data sets for normal distribution, using the Shapiro-Wilk test. If normal distribution was proved, unpaired t-test was used for comparison and if data did not have a normal distribution comparisons were performed using the Mann-Whitney test. In all behavioral analysis (OF, MM, GR, RI and NOR) we first tested the datasets for normal distribution, using the Shapiro-Wilk test and identified outliers. After normal distribution was probed and outliers excluded, unpaired t-test was used to compare the two groups. In all cases, differences were considered significant when *P* values were smaller than 0.05.

## ACKNOWLEDGMENTS

We thank Gabriel Anessetti (Dpto. de Histología y Embriología, Facultad de Medicina, UdelaR) for help with the immunofluorescence of renal tissues, Lucilla Pizzo for her advice on the manuscript and Paola Lepanto for her help with image processing.

## Competing Interests

None declared

## FUNDING

This study was supported by: Programa para el Desarrollo de las Ciencias Básicas (PEDECIBA) and Sistema Nacional de Investigadores-Agenica Nacional de Investigación e Innovación (ANII) to MF, MC-R, MB, VP-E, MC, PL, NL, CE, FI and JLB; CSIC I+D 2016-557 (Universidad de la República, Uruguay) to FI and JLB; Comisión Académica de Posgrados (CAP-UdelaR) to MF; FOCEM – Fondo para la Convergencia Estructural del Mercosur (COF 03/11). Santhosh Girirajan is supported by NIH R01 GM121907 and Corrine Smolen is supported by NIH T32-GM102057.

## Supplementary Figure Legends

**Supplementary S1 Fig:** Nanopore sequence alignments to the genomic *Ccdc28b* region.

**Supplementary S2 Fig:** Full length western blot gels assessing the levels of Ccdc28b in brain and muscle.

**Supplementary S3 Fig:** Confocal images showing A) amygdala (bar= 50 μm), B) hippocampal CA1 (bar= 50 μm), C) dentate gyrus (bar= 50 μm). DAPI (magenta) and anti-ACIII antibody (green) were used for nucleus and cilia visualization respectively.

**Supplementary S4 Fig:** Comparison of basal glycemia at three timepoints during the HFD experiment for each genotype (*Ccdc28b mut* and wt).

**Supplementary S5 Fig:** Comparison of average phenotypic scores of probands with *CCDC28B* mutations with a distribution of average scores of 7 or 8 probands drawn randomly 1000 times. Note that the phenotypic data were available for all eight or seven probands for the tested phenotypes.

## Notes

### Competing Interest Statement

The authors have declared no competing interest.

